# Loss of transgene expression limits liver gene therapy in primates

**DOI:** 10.1101/2022.03.24.485675

**Authors:** Jenny A. Greig, Camilo Breton, Kelly M. Martins, Yanqing Zhu, Zhenning He, John White, Peter Bell, Lili Wang, James M. Wilson

**Author notes:** Corresponding author: James M. Wilson, MD, PhD, Gene Therapy Program, Perelman School of Medicine, University of Pennsylvania, 125 South 31^st^ Street, Suite 1200, Philadelphia, PA 19104, USA.

## Abstract

Efforts to improve liver gene therapy have focused on next-generation adeno-associated virus (AAV) vector capsids, transgene delivery, and immunomodulating drugs, such as corticosteroids, to avoid destructive T-cell responses. We conducted a detailed characterization of AAV transduction in nonhuman primate liver across multiple capsids and transgenes to better define interactions that may limit stable and efficient transgene expression. We show that the initial transduction of hepatocytes is high, but the transduction rapidly diminishes to a lower stable baseline of <1% of cells, even though ~10% of the cells retain vector DNA that is localized within the nucleus. Further characterization showed genomic vector integration at frequencies of 1/100 to 1/1,000 genomes, suggesting that one mechanism for stable expression may occur via genome integration. These studies suggest that strategies to enhance durable transgene expression by activating retained nuclear episomes or by increasing the frequency of vector integrations may improve liver directed AAV gene therapy.

## Main Text

The goal of our studies was to define the mechanisms that limit efficient and durable transgene expression following liver gene therapy with AAV vectors. Previous studies with complete preclinical and clinical datasets suggest that nonhuman primates (NHPs) are better suited for evaluating key aspects of vector performance than other animal models^1–4^. We conducted initial studies in rhesus macaques using macaque-derived β-choriogonadotropic hormone (rh-β-CG) as the transgene. rh-β-CG is secreted and has a short half-life in serum, allowing for longitudinal, real-time readouts of transgene transcription. As rh-β-CG should be viewed as a self-protein in macaques, confounding adaptive immune responses to the transgene product or to transgene-expressing cells are unlikely. We evaluated two clade E capsids that have been used in multiple clinical trials, AAV8 and AAVrh10 (N = 6 NHPs/vector). As an essential and unique aspect of our studies, we analyzed three sequential liver tissue samplings from each animal, including biopsies at days 14 and 98 and necropsy at day 182. These analyses included serum rh-β-CG, liver transgene DNA and RNA by quantitative polymerase chain reaction (qPCR), and cellular distribution of DNA and rh-β-CG protein expression by *in situ* hybridization (ISH) and immunohistochemistry (IHC), respectively.

We observed similar levels and profiles of rh-β-CG protein expression in serum for each capsid; peak levels were achieved by day 7, followed by a gradual decline to stable levels 3- to 6-fold lower than the peak (Fig. 1a) without transaminase elevations (Fig. 1b). Statistically significant reductions in total vector DNA and RNA occurred over time, although the reductions were greater in RNA than in DNA (Fig. 1c,d). While DNA levels decreased further over the two later time points, RNA levels were stable between day 96 and 182 after the initial decline relative to day 14 (Fig. 1c,d).

**Figure 1.**
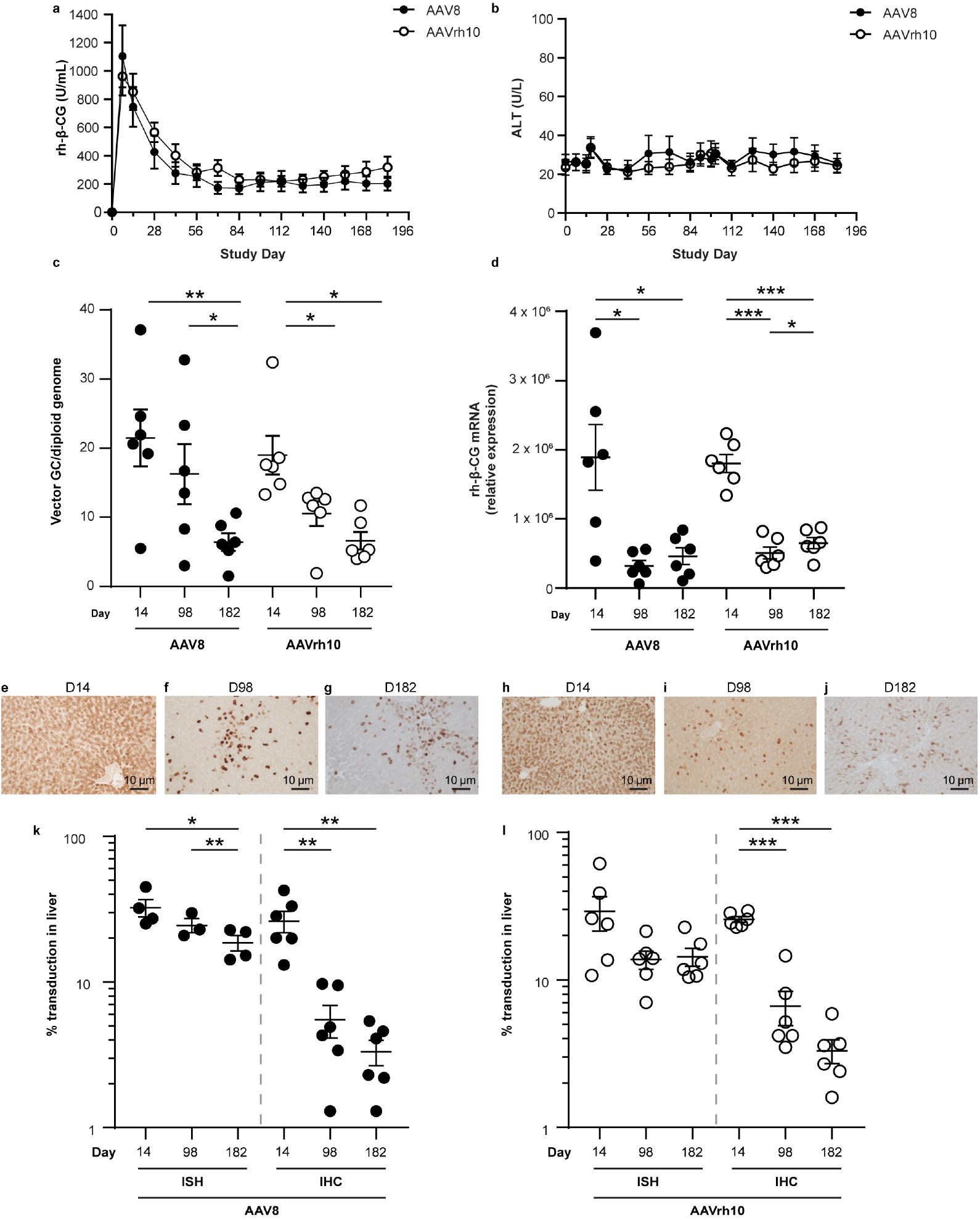
Initial peak followed by a decline to lower stable levels of self-transgene after IV administration of AAV vectors to NHPs. NHPs received intravenous (IV) injections of 10^13^ GC/kg of AAV8 or AAVrh10 vectors expressing the self-transgene rh-β-CG (N = 6/group). Serum rh-β-CG levels were evaluated throughout the in-life phase for transgene expression (a) and alanine aminotransferase (ALT) (b). Liver tissue was harvested during a liver biopsy procedure (14- or 98-days post-vector administration) or at the time of necropsy (182-days post-vector administration). Liver tissue was harvested via liver biopsy (14- or 98-days post-vector administration) or necropsy (182 days post-vector administration). (c) DNA and (d) RNA were extracted from liver samples to evaluate the number of vector GCs and transgene RNA levels, respectively. IHC for the rh-β-CG transgene was performed on liver samples (brown staining) for animals administered with AAV8 (e-g) or AAVrh10 (h-j). ISH was performed on liver samples using the RNAscope Multiplex Assay. Hybridized probes were imaged with a fluorescence microscope. IHC was performed for the rh-β-CG protein. The vector DNA was quantified by ISH and transgene expression by IHC as the percentage of AAV-positive cells for NHPs administered with AAV8 (k) or AAVrh10 (l). Values are presented as the mean ± SEM. *p < 0.05, **p < 0.01, ***p < 0.001.

To elucidate the mechanism governing this rapid decline in expression, we evaluated the cellular distribution of rh-β-CG protein expression by IHC (Fig. 1e–g for AAV8 and 1h–j AAVrh10; Supplementary Fig. S1) and the cellular distribution of nuclear DNA by ISH using a probe specific for vector DNA as part of a probe pair to target DNA and RNA separately as described in Methods (Fig. 1k for AAV8 and 1l AAVrh10). The number of rh-β-CG-positive cells declined 3- to 5-fold from day 14 to day 98 and then remained relatively stable through day 182, consistent with the kinetics of rh-β-CG in serum and transgene RNA in liver. The ~4-fold reduction in rh-β-CG-expressing cells was not associated with a commensurate reduction in DNA-containing cells, which decreased by only 24%–53%. A more detailed discussion of the pattern of DNA localization is provided below.

We next evaluated the same parameters of gene transfer and expression in macaques infused with AAV8 vectors expressing one of three transgenes, including the reporter gene enhanced green fluorescent protein (eGFP) and the human and macaque versions of low-density lipoprotein receptor (hLDLR and rhLDLR, respectively; N = 2/transgene). These studies allowed us to assess the role of adaptive immunity in the efficiency and stability of transgene expression within a range comprising the highly immunogenic protein eGFP to the non-immunogenic self-protein rhLDLR.

Throughout the in-life phase of the study, we utilized serum low-density lipoprotein (LDL) levels as an indirect assessment of transgene expression (Fig. 2a–c). Animals that received the rhLDLR vector showed an acute and substantial reduction in serum LDL, which returned to levels close to or at baseline within 30 days (Fig. 2a). We observed a similar pattern of serum LDL for the hLDLR vector, but with a lower magnitude of transient reduction (Fig. 2b). As expected, we did not observe a reduction in serum LDL for the eGFP vector (Fig. 2c). Serum transaminases tracked with immunogenicity, ranging from no elevations with rhLDLR (Fig. 2d) to mild elevations with hLDLR (Fig. 2e), to a sharp and transient increase with eGFP (Fig. 2f).

**Figure 2.**
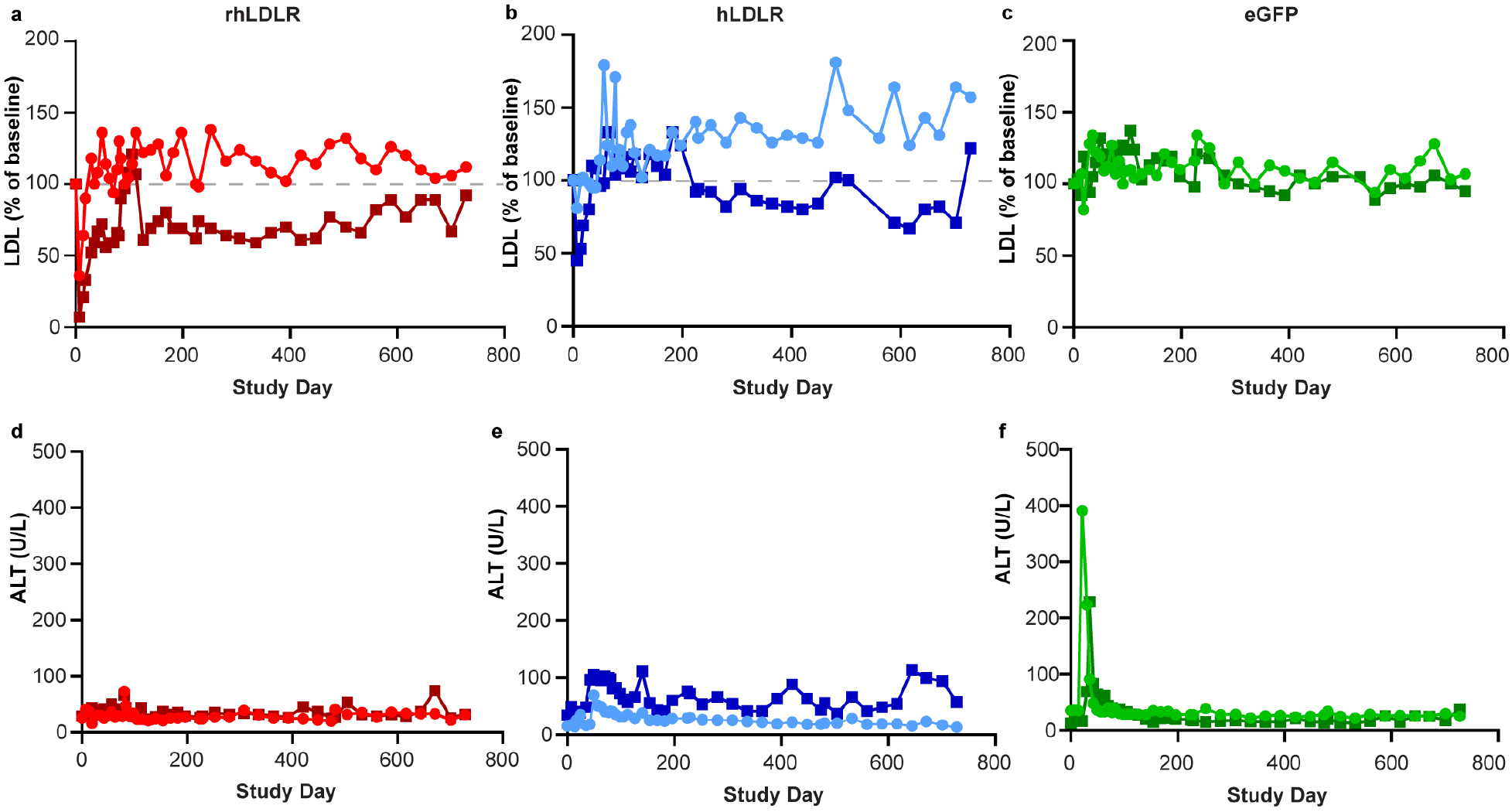
Comparison of the duration of expression of self, human, and non-self transgenes following IV administration of AAV vectors. NHPs received IV injections of 10^13^ GC/kg of AAV8 vectors expressing rhLDLR), hLDLR, or eGFP (N = 2/group). Serum LDL (a-c) and ALT (d-f) levels were evaluated throughout the in-life phase.

We performed tissues studies as previously described for the rh-β-CG experiments, although four samples were available for each animal (biopsies at days 14, 77, and 224 and necropsy at day 760). Analyses of vector DNA and transgene RNA, as measured by qPCR, followed the same trend over time as that for rh-β-CG across all transgenes, showing reductions in both DNA (Fig. 3a) and RNA (Fig. 3b), with greater losses in RNA compared with DNA. The extent of these reductions was much greater for the eGFP transgene, with DNA diminishing >3,000-fold and RNA >55,000-fold, compared with rhLDLR, with DNA diminishing 6-fold and RNA 29-fold. The hLDLR vector results were more similar to the data for the rhLDLR vector, showing DNA and RNA reductions of 9-fold and 227-fold, respectively.

**Figure 3.**
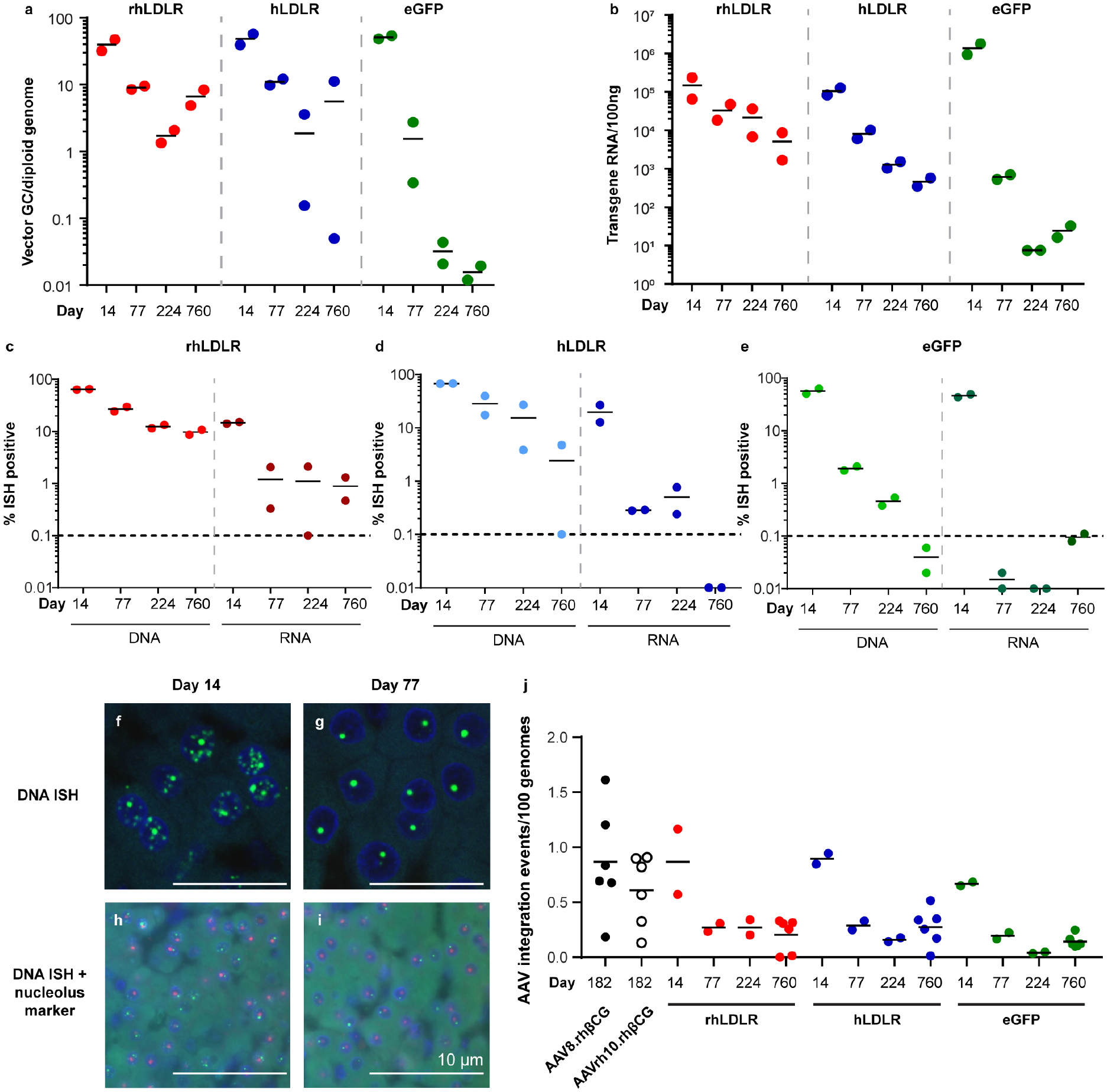
Quantification and localization of vector DNA following IV administration of AAV vectors to NHPs. NHPs received IV injections of 10^13^ GC/kg of AAV8 vectors expressing rhLDLR, hLDLR, or eGFP (N = 2/group). Liver tissue was harvested during a liver biopsy (14-, 77-, and 224-days post-vector administration) or necropsy (760-days post-vector administration). (a) DNA and (b) RNA were extracted from liver samples to evaluate the number of vector GCs and transgene RNA levels, respectively. ISH was performed on liver samples using the RNAscope Multiplex Assay. The probes used were non-overlapping probe pairs, where one probe is specific for DNA (binding to the anti-sense strand) and the second probe hybridizes to RNA. Hybridized probes were imaged with a fluorescence microscope. The vector DNA and transgene RNA were quantified as the percentage of AAV-positive cells for animals administered with AAV vectors expressing rhLDLR (c), hLDLR (d), or eGFP (e). The dashed lines indicate the level where background can be observed due to autofluorescence. Hybridized probes were imaged on some sections with a confocal microscope (f, g). Green, vector DNA; blue, DAPI (nuclear counterstain). Sections were also co-stained with a nucleolus marker (fibrillarin antibody) shown in red (h, i). (j) The number of AAV integration loci in all injected NHPs was determined by ITR-seq. The number of integration events per 100 genomes is equivalent to the percentage of cells in the liver with AAV integration (one liver sample per NHP was evaluated, except for samples harvested at day 760 where three sites per NHP were evaluated). ISH for vector DNA was performed on liver samples from NHPs administered the rhLDLR vector using the RNAscope Multiplex Assay. Values are presented as the mean. *p < 0.05, **p < 0.01, ***p < 0.001.

We performed ISH to characterize the number of cells harboring intranuclear vector DNA in comparison to those with cytoplasmic transgene-derived RNA (Supplementary Figs. S2 and S3). The pattern observed for the rhLDLR vector was similar to the rh-β-CG vector results, with high numbers of DNA- and RNA-expressing cells at day 14 (65% and 15%, respectively) followed by a 7-fold reduction of DNA over 2 years and a 12-fold reduction in RNA-expressing cells by day 77. The number of RNA-expressing cells then remained stable through day 760, reaching a steady state of ~1% RNA-expressing cells (Fig. 3c; Supplementary Figs. S2 and S3). The pattern for hLDLR was essentially the same, although with greater reductions in DNA (28-fold) and RNA (~3,000-fold), with RNA-expressing cells falling below the 0.1% threshold of detection at day 760 (Fig. 3d; Supplementary Figs. S2 and S3). Animals that received the eGFP vector exhibited the same high level of gene transfer and transgene expression based on analyses for day 14, although the expression dropped to undetectable levels by day 760 for DNA and day 77 for RNA (Fig. 3e; Supplementary Figs. S2 and S3).

The presence of many non-expressing hepatocytes that continued to harbor vector DNA compelled us to more fully characterize the structure and location of the vector genomes detected by ISH. Confocal images of tissues subjected to ISH with a DNA-specific probe to detect vector DNA as part of a probe pair to target DNA and RNA separately (as described in Methods) revealed two patterns of intranuclear hybridization at day 14: a diffuse granular pattern with a single bright circular structure within some nuclei (Fig. 3f). Evaluations at later time points demonstrated retention of the circular structures, with a loss of the background granular staining (Fig. 3g). We analyzed additional tissue sections for vector DNA by ISH and for fibrillarin as a nucleolus marker by IHC. There was no overlap between these two structures, indicating that the single nuclear domain that harbors vector genomes is not related to the nucleolus (Fig. 3h,i).

We and others have shown integration of AAV genome sequences into chromosomal DNA in the setting of DNA repair following double-stranded breaks^5–11^. We evaluated liver DNA from the rh-β-CG- and LDLR/eGFP-treated animals for integrated vector sequences using a technique called inverted terminal repeat sequencing (ITR-seq)^5^. This method captures chromosomal sequences flanking insertion sites through PCR amplification of sheared DNA ligated with adapters using oligonucleotides specific to the AAV ITR and the linkers. Analysis of day-182 tissues from rh-β-CG-vector-treated animals showed integration events at frequencies of 1/100 to 1/1,000 genomes (Fig. 3j). A time course of integrations performed in the eGFP/LDLR study showed similar levels of integration events, which declined between days 14 and 77 and subsequently stabilized to levels of 0.01–0.52 AAV integration events per 100 genomes (Fig. 3j). These integrations occurred randomly throughout the genome, with a bias in and around genes that are highly expressed in the liver (Supplementary Fig. S4).

Our studies in NHPs provide a model to clarify why some clinical trials of liver gene therapy have been underwhelming in terms of efficiency and durability. Our model indicates that the issue is more strongly related to sustained transgene expression, rather than vector genome delivery. Modest doses of vector can achieve expression in most hepatocytes, although in primates, in contrast to mice, expression is lost in most cells between 2 and 6 weeks. This loss of expression occurs with self, non-immunogenic transgene products in the absence of overt evidence of adaptive immunity or inflammation, such as what we observed in our study of gene therapy in a 12-year-old patient with Crigler-Najjar type I, a condition associated with severe hyperbilirubinemia due to a deficiency of uridine diphosphate glucuronosyl transferase 1A1 (UGT1A1). Soon after vector infusion, this patient’s bilirubin levels, which were determined as a real-time readout of hepatic *UGT1A1* expression, rapidly decreased to normal, followed by a return to high pretreatment levels over the subsequent 4–6 weeks without evidence of liver inflammation^12^. However, our studies do demonstrate how inflammation associated with non-self transgenes, as evidenced by transaminitis, can lead to more substantial reductions in vector DNA and transgene expression. This finding is consistent with studies in hemophilia gene therapy, which have shown an association between transaminase elevations and reductions in circulating transgene-derived clotting factors^13^, but in the absence of a transgene-related immune response.

The presence of large persistent intranuclear domains of AAV genomes in our AAV gene therapy studies is reminiscent of the replication centers observed during viral infection with adenovirus and herpes simplex virus. One hypothesis is that transcriptionally active genomes located within these structures, or elsewhere in the nucleus, undergo epigenetic silencing, similar to how intranuclear genomes of viruses such as herpesviruses are inactivated. Another potential explanation is that genomes that are transcriptionally active soon after delivery are unstable^14^, such as the transiently appearing vector DNA that is widely dispersed in the nucleus, while the vector genomes within the persisting discrete nuclear domains were never active. Alternatively, these internuclear foci could result from the slow conversion of circular monomeric episomes that occurs after vector delivery into multimeric AAV genome concatemers, as previously described in skeletal muscle^15^. These concatemeric structures would provide many binding sites for ISH probes, resulting in a bright localized signal, and may not allow efficient transgene expression.

The demonstration of integrated vector genomes at levels of 1/100 to 1/1,000 genomes inform the model in two ways. The fact that integration events are far less common (<2%) than the number of cells that express at day 14 (i.e., >20%) and the number of cells that retain vector DNA within the nuclear domain (i.e., ~10%) indicates that episomal forms of the vector genome must be responsible for early transcription and the formation of discrete nuclear domains. Integrated vector sequences are detected at overall frequencies of <1% at long term timepoints. However, this integration occurs at the same low frequency as the number of cells that stably express non-foreign transgenes, suggesting that stable expression may be the result of integrated genomes. The durability of residual transgene expression observed following gene therapy in large animal models and humans with liver diseases such as hemophilia B is also consistent with integration, as a non-replicating episome should have been diluted via normal hepatocyte turnover. Validation of this hypothesis would require evidence that most of these integration events contain transcriptionally active intact minigene cassettes and would require a comparison of self and non-self transgenes.

Our findings have implications for improving the efficacy of liver gene therapy. One strategy is to enable the retained episomal AAV genomes located in the discrete nuclear domain to be more transcriptionally active by incorporating cis elements, such as different promoters and insulators, or by using drugs that prevent epigenetic silencing. This approach may be effective regardless of whether these vector genomes were active and then silenced or simply were never active. Another strategy, based on the proposal that integrated genomes are more likely to confer durable expression, would be to increase the number of vector integrations into safe harbor sites, as is the goal of some editing strategies for treating loss-of-function diseases. In this scenario, we can view editing-based gene insertions as an extension of gene therapy, in which the editing-based insertions are directed to a specific chromosomal location with higher frequency than the random background of insertions observed following gene therapy.

## Methods

### Data availability statement

All data discussed in the manuscript are available in the main text or supplementary materials. Complete clinical pathology data can be obtained upon request.

### AAV vector production

All adeno-associated virus (AAV) vectors were produced by the Penn Vector Core at the University of Pennsylvania as previously described^16^. Briefly, plasmids expressing rhesus choriogonadotropin (rh-β-CG), codon-optimized rhesus low-density lipoprotein receptor (rhLDLR), codon-optimized human low-density lipoprotein receptor (hLDLR), and enhanced green fluorescent protein (eGFP) from the thyroxine-binding globulin (TBG) promoter were packaged within the AAV8 capsid. A vector expressing rh-β-CG from the TBG was also packaged with the AAVrh10 capsid. Unique biological materials are available upon request, pending permission by the authors and/or patent holder(s).

### Animal studies

The Institutional Animal Care and Use Committee of the University of Pennsylvania approved all animal procedures in this study. We obtained wild-type rhesus macaques aged 3–6 years (N = 18) from Covance Research Products, Inc. (Alice, TX). We conducted nonhuman primate (NHP) studies at the University of Pennsylvania within a facility that is registered with the United States Department of Agriculture, accredited by the American Association for Accreditation of Laboratory Animal Care, and assured by the Public Health Service. As previously described^17^, we housed animals in stainless steel squeeze-back cages with perches. All cage sizes and housing conditions were compliant with the Guide for the Care and Use of Laboratory Animals. A 12-h light/dark cycle was maintained and controlled via an Edstrom Watchdog system. Animals were fed Certified Primate Diet 5048 (PMI Feeds Inc., Brentwood, MO, USA) two times per day (morning and evening). We also provided an additional variety of food treats that were fit for human consumption, including fruits, vegetables, nuts, and cereals, daily as part of the standard enrichment process. Manipulanda such as kongs, mirrors, a puzzle feeder, and raisin balls were provided daily. Animals also received visual enrichment and daily human interaction. All interventions were performed during the light cycle, and animals were fasted overnight prior to being anesthetized.

On study day 0, rhesus macaques received 10^13^ genome copies (GC)/kg of AAV8.TBG.rh-β-CG (N = 6), AAVrh10.TBG.rh-β-CG (N = 6), AAV8.TBG.rhLDLR (N = 2), AAV8.TBG.hLDLR (N = 2), or AAV8.TBG.eGFP (N = 2) as a 10-mL infusion of vector into the saphenous vein at a rate of 1 mL/min via an infusion pump (Harvard Apparatus, Holliston, MA). Macaques were anesthetized and blood was collected on selected days via the femoral vein for analysis of complete blood counts, clinical chemistries, and coagulation panels by Antech (Irvine, CA). We determined neutralizing antibody (NAb) titers using serum samples taken prior to the initiation of the study, as previously described^18^. All animals had NAb titers of <1/5 for the administered AAV capsid prior to vector administration. At baseline and throughout the in-life phase of the study, we evaluated the animals for serum biomarkers to determine the transgene expression of either rh-β-CG by enzyme-linked immunosorbent assay as previously described^19^ or low-density lipoprotein (LDL) levels as a biomarker for rhLDLR and hLDLR transgene expression. Lipid panel analysis was performed by Antech GLP (Morrisville, NC).

We performed liver biopsies on NHPs throughout the in-life phase of the studies (on study days 14 and 98 for NHPs receiving AAV8.TBG.rh-β-CG and AAVrh10.TBG.rh-β-CG and on study days 14, 77, and 224 for NHPs receiving AAV8.TBG.rhLDLR, AAV8.TBG.hLDLR, or AAV8.TBG.eGFP). We conducted a mini-laparotomy procedure to isolate liver tissue. We divided the collected samples for histopathology analysis (i.e., fixed in 10% neutral buffered formalin) and molecular analysis (i.e., frozen on dry ice and stored at −60°C or colder).

### Necropsy

At day 182 post-vector administration for NHPs administered with rh-β-CG vectors and at day 760 for animals administered rhLDLR, hLDLR, and eGFP vectors, rhesus macaques were euthanized and necropsied.

### Vector genome copy and transgene RNA analysis

We snap-froze tissue samples at the time of biopsy or necropsy and extracted DNA using the QIAamp DNA Mini Kit (Qiagen, Valencia, CA). We isolated DNase-treated total RNA from 100 mg of tissue. RNA was quantified by spectrophotometry and reverse transcribed to cDNA using random primers. We detected and quantified vector GC levels in extracted DNA and transgene expression in extracted RNA using real-time polymerase chain reaction (PCR), as previously described^16, 17^. Briefly, vector GC and RNA levels were quantified using primers and a probe designed for a vector-specific sequence.

### Immunohistochemistry for CG

We performed immunohistochemistry (IHC) for CG on formalin-fixed, paraffin-embedded liver sections. The sections were deparaffinized through a xylene and ethanol series, boiled for 6 min in 10 mM citrate buffer (pH 6.0) for antigen retrieval, and treated with 2% H_2_O_2_ (15 min), avidin/biotin-blocking reagents (15 min each; Vector Laboratories), and blocking buffer (1% donkey serum in phosphate-buffered saline [PBS] + 0.2% Triton for 10 min). We then incubated the sections with a rabbit serum against human CG (Abcam ab9376, diluted 1:200) for 1 h at 37°C followed by biotinylated secondary antibodies (45 min at room temperature; Jackson ImmunoResearch, West Grove, PA) diluted in blocking buffer according to the manufacturer’s recommendations. We employed a Vectastain Elite ABC kit (Vector Laboratories) with 3,3’-diaminobenzidine as a substrate to stain bound antibodies according to the manufacturer’s instructions. No counterstain was applied to the sections to facilitate quantification.

For quantification, we acquired ten random images from each IHC-stained section with a 10x objective. Using ImageJ software (Rasband, W. S., National Institutes of Health, USA; http://rsb.info.nih.gov/ij/), we measured the area positive for CG IHC and the area occupied by central and portal veins for each image, which was then used to calculate the average percentage of CG-positive area of liver tissue while excluding vein areas.

### *In situ* hybridization for vector DNA and transgene RNA

We performed *in situ* hybridization (ISH) on formalin-fixed paraffin-embedded liver sections using the RNAscope Multiplex Fluorescent Reagent Kit v2 Assay (Advanced Cell Diagnostics, ACD) following the manufacturer’s protocol. Probes for two-plex ISH were synthesized by ACD and designed as non-overlapping probe pairs, where one probe is specific for DNA (binding to the anti-sense strand) and the second probe hybridizes to RNA and to the sense DNA strand. Probes for DNA were stained first and detected by Opal 520 precipitates (imaged with a filter set for fluorescein isothiocyanate), and probes binding to RNA/DNA were stained in a second step with Opal 570 deposits (imaged with a rhodamine filter set). Reactive Opal dyes were purchased from Akoya Biosciences, and images were taken with a Nikon Eclipse Ti-E fluorescence microscope. For some sections, we acquired confocal images using a Leica TCS SP5 confocal microscope with an acousto-optical beam splitter.

To quantify ISH-positive cells, we scanned stained sections with an Aperio Versa fluorescence slide scanner (Leica Biosystems) and analyzed the sections using Visiopharm software (Hoersholm, Denmark) with apps that detect either the probe signal for DNA inside 4′,6-diamidino-2-phenylindole (DAPI)-stained nuclei or the probe signal for RNA in the cytoplasm.

### ISH with nucleolus localization

After performing ISH as described above, we further stained some sections with an antibody against fibrillarin as a nucleolar marker. After treatment with 0.5% Triton/PBS for 2 h and blocking with 1% donkey serum/0.2% Triton in PBS for 1 h, the sections were incubated with a rabbit anti-fibrillarin antibody (Abcam ab166630, diluted 1:100 in 0.5% Triton/PBS) overnight at 4°C, followed by Cy5-labeled secondary antibody (donkey anti-rabbit from Jackson ImmunoResearch, 1:100 in 0.5% Triton/PBS, 1 h). We mounted sections using ProLong Gold Antifade Mountant with DAPI (Invitrogen).

### Next-generation sequencing for AAV integration analysis

To evaluate AAV genomic integrations, we performed ITR-seq^5^. Briefly, we performed an unbiased genome-wide detection of off-targets on purified liver DNA that was sheared using an ME220 focused ultrasonicator, end-repaired, A-tailed, and ligated to special adapters, as described for AMP-Seq. Using AAV-ITR and adapter-specific primers, we amplified ITR-containing DNA fragments and generated next-generation-sequencing-compatible libraries. We sequenced the DNA on a MiSeq instrument and mapped the obtained reads to the rhesus macaque reference genome and the administered AAV vector genome. We used a custom script to identify AAV integration sites from the mapped reads, as previously described^5^. The genomic location of AAV integration loci were determined by ITR-seq and annotated as being within a gene-coding region (genic) or outside of a gene-coding region (intergenic) according to the rhesus macaque RefSeqGene annotation. Integrations within genic regions were further annotated by the distribution of RNA expression in the human liver (data taken from the Human Genome Atlas [www.proteinatlas.org]). Expression levels were determined by normalized expression (nx) in the human liver for each annotated gene. Categories were determined as follows: genes not expressed in liver: 1 < nx; genes with low expression in liver: 1 ≥ nx < 10; genes with medium expression in liver: 10 < nx > 100; genes with high expression in liver: 100 ≥ nx. A random distribution represents 10,000 randomly computationally generated genomic loci in the rhesus macaque genome. The number of AAV integration events in each gene category is presented as a fold-change over random sequences. A value of 1 would represent no difference from the distribution of random loci.

### Statistical analysis

Comparisons between time points were performed for vector GC, transgene RNA, DNA ISH, and RNA ISH levels using paired *t*-tests in the “t.test” function within the R Program (version 4.0.0). We conducted comparisons between vector GCs and transgene RNA and quantifications of DNA ISH and IHC using linear mixed-effect modeling using the “lme” function in the “nlme” package for R. A p-value of <0.05 was considered significant.

## Acknowledgments

We thank Alexa N. Avitto, Joanna K. Chorazeczewski, Rucha Fadnavis, Thomas Furmanak, Matthew Jennis, and Melanie K. Smith for their support of this project and instrumental contributions. We also thank Jing-Xu Zhu for confocal imaging, nucleolus co-staining, and help with ISH, and Tianying Jiang for scanning the slides used for quantification. We thank Nathan Denton for assistance with manuscript preparation and graphics, and Hanying Yan for help with statistical analysis. We thank the Biostatistics and Bioinformatics Core, Immunology Core, Nucleic Acid Technologies Core, Pathology Core, Program for Comparative Medicine, and Vector Core of the Gene Therapy Program at the University of Pennsylvania for study support. All vectors were produced by the Penn Vector Core.

## Author Contributions

J.A.G. – conceptualization; data curation; formal analysis; investigation; methodology; supervision; visualization; writing-original; writing-review and edits

C.B. – conceptualization; data curation; formal analysis; investigation; methodology; supervision; visualization; writing-original; writing-review and edits

K.M.M. – formal analysis; investigation; methodology; visualization; writing-review and edits

Y.Z. – methodology

Z.H. – data curation; formal analysis

J.W. – data curation; formal analysis

P.B. – data curation; formal analysis; investigation; methodology; software; supervision

L.W. – conceptualization; data curation; formal analysis; methodology; supervision; writing-review and edits

J.M.W. – conceptualization; funding acquisition; investigation; methodology; supervision; writing-original; writing-review and edits

## Supplementary Data

**Figure S1.**
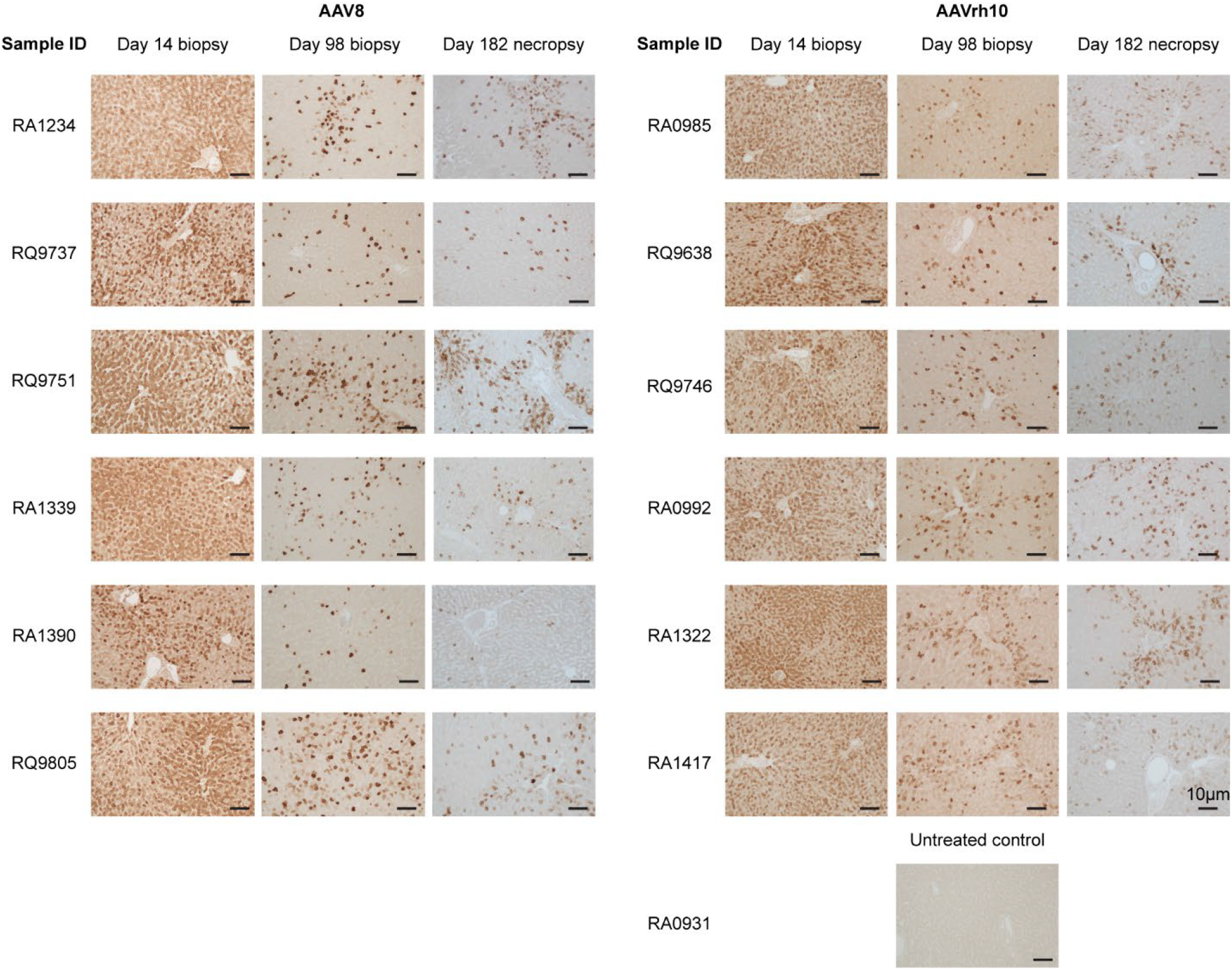
Reduction in self-transgene transduced hepatocytes over time following IV administration of AAV vectors to NHPs. NHPs received IV injections of 10^13^ GC/kg of AAV8 or AAVrh10 vectors expressing the self-transgene rh-β-CG (N=6 per group). Liver tissue was harvested during a liver biopsy procedure (14- or 98-days post-vector administration) or at the time of necropsy (182 days post-vector administration). IHC was performed on liver samples for the CG transgene (brown staining).

**Figure S2.**
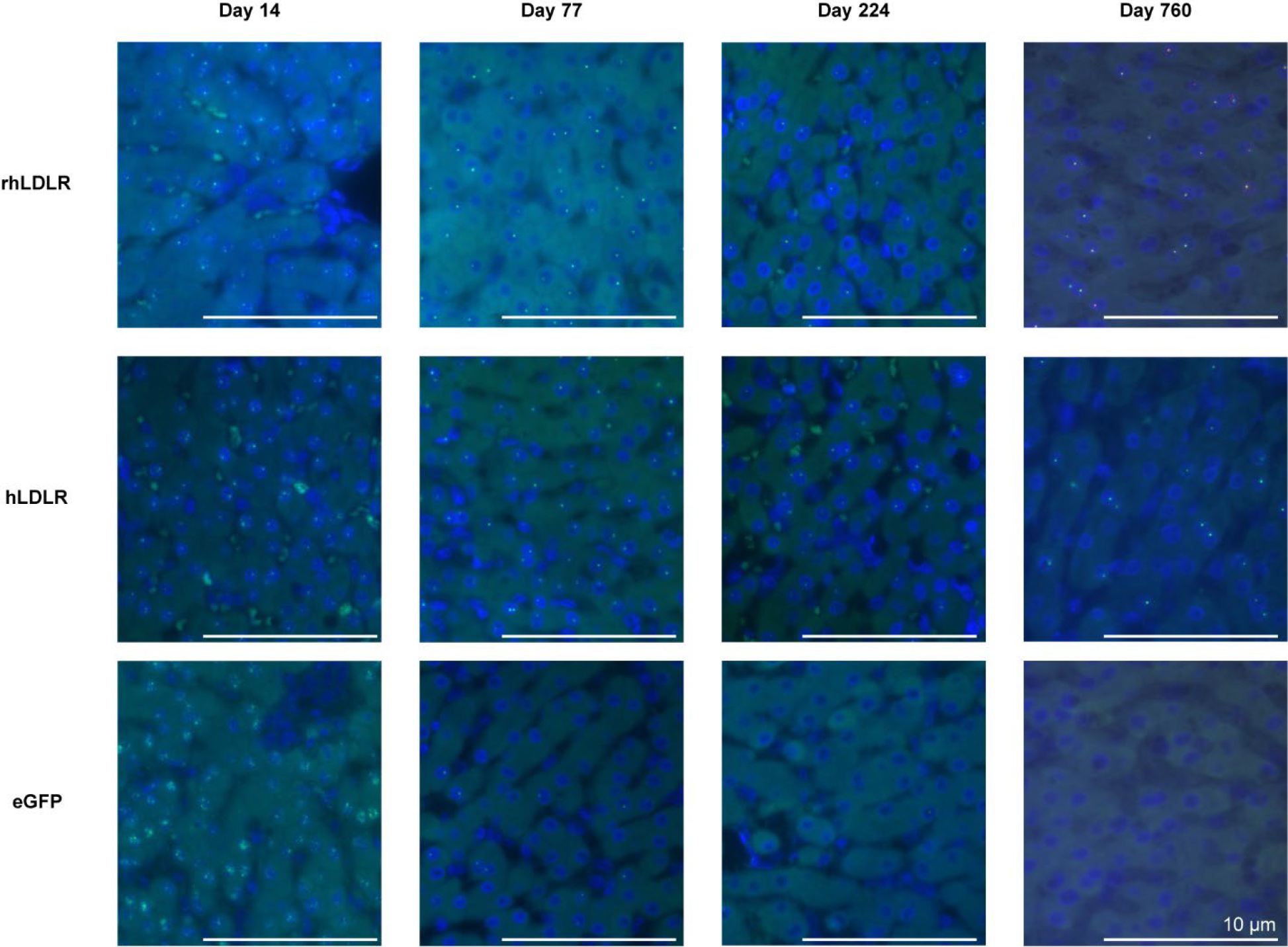
Loss of vector DNA correlates with difference from self following IV administration of AAV vectors. NHPs received IV injections of 10^13^ GC/kg of AAV8 vectors expressing rhLDLR, hLDLR, or eGFP (N=2/group). Liver tissue was harvested during a liver biopsy procedure (14-, 77-, and 224-days post-vector administration) or at the time of necropsy (760 days post-vector administration). ISH was performed on liver samples using the RNAscope Multiplex Assay. The probes used were non-overlapping probe pairs, where one probe is specific for DNA (binding to the anti-sense strand). Hybridized probes were imaged with a fluorescence microscope. Green, vector DNA; blue, DAPI (nuclear counterstain).

**Figure S3.**
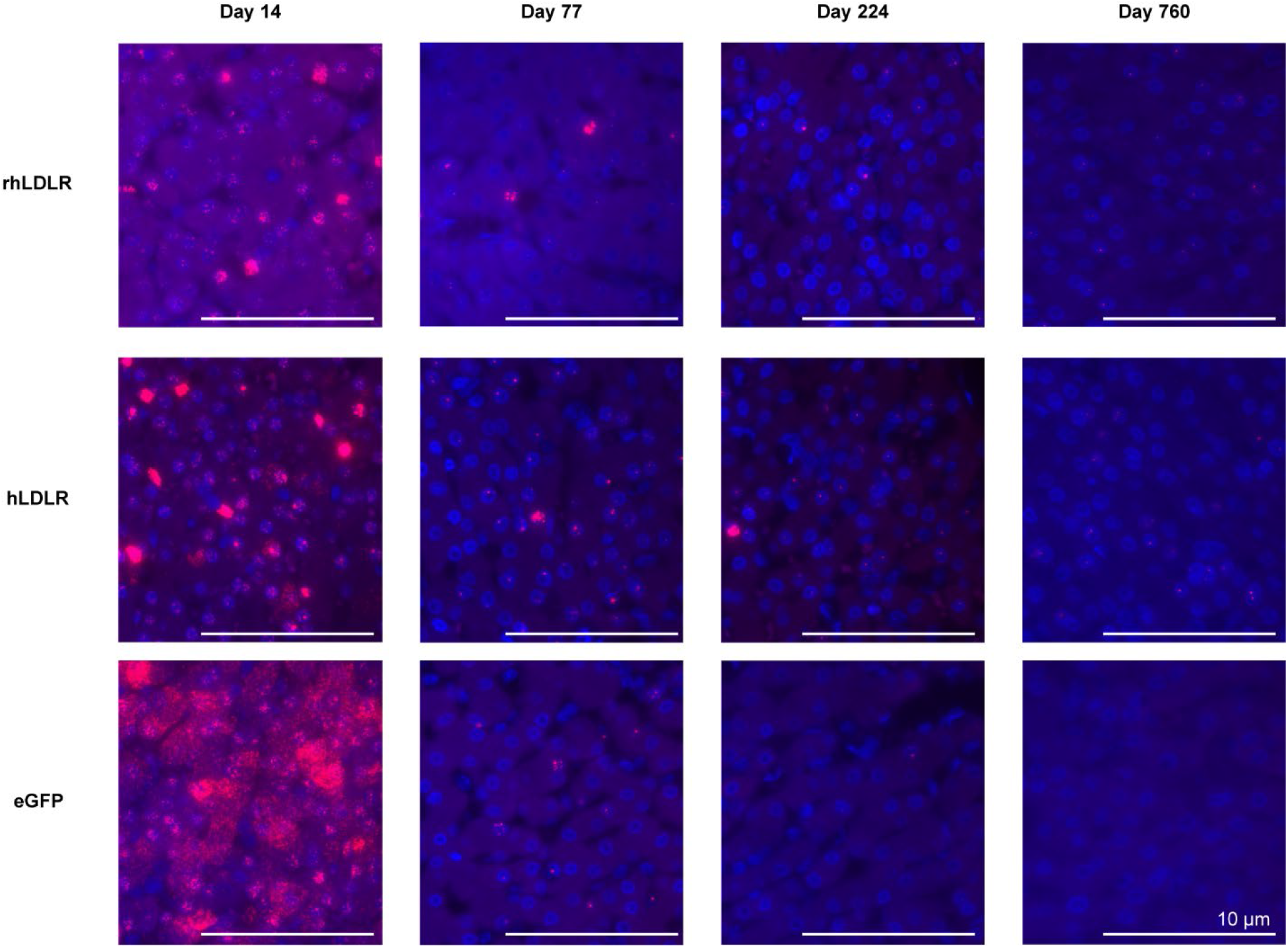
Loss of vector RNA correlates with difference from self following IV administration of AAV vectors. NHPs received IV injections of 10^13^ GC/kg of AAV8 vectors expressing rhLDLR, hLDLR, or eGFP (N=2/group). Liver tissue was harvested during a liver biopsy procedure (14-, 77-, and 224 days post-vector administration) or at the time of necropsy (760 days post-vector administration). ISH was performed on liver samples from NHPs using the RNAscope Multiplex Assay. Probes hybridized to RNA were imaged with a fluorescence microscope. Red, transgene RNA; blue, DAPI (nuclear counterstain).

**Figure S4.**
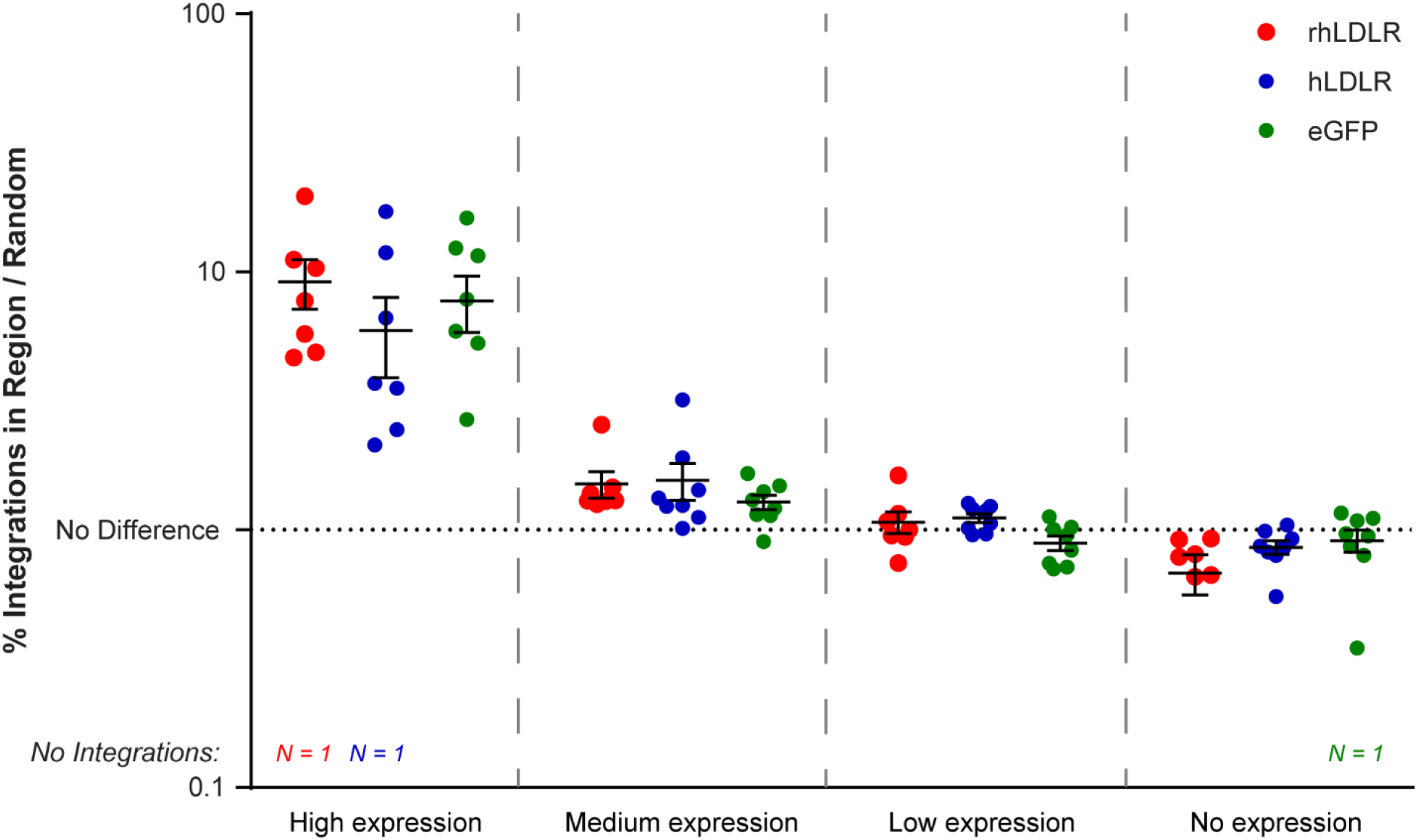
Preferential integration of the AAV genome into active genes in the liver of NHPs following IV administration of AAV vectors. NHPs received IV injections of 10^13^ GC/kg of AAV8 vectors expressing rhLDLR, hLDLR, or eGFP (N=2/group). Liver tissue was harvested during a liver biopsy procedure (14-, 77-, and 224-days post-vector administration) or at the time of necropsy (760-days post-vector administration). The genomic location of AAV integration loci were determined by ITR-seq and annotated as being within a gene-coding region (genic) or outside of a gene-coding region (intergenic) according to the rhesus macaque RefSeqGene annotation. Integrations within genic regions were further annotated by the distribution of RNA expression in the human liver (data taken from the Human Genome Atlas [www.proteinatlas.org]). Expression levels were determined by normalized expression (nx) in the human liver for each annotated gene. Categories were determined as follows: genes not expressed in liver: 1 < nx; genes with low expression in liver: 1 ≥ nx < 10; genes with medium expression in liver: 10 < nx > 100; genes with high expression in liver: 100 ≥ nx. A random distribution represents 10,000 randomly computationally generated genomic loci in the rhesus macaque genome. The number of AAV integration events in each gene category is presented as a fold-change over random sequences. A value of 1 would represent no difference from the distribution of random loci. Data presented as mean ± SEM.

